# Itaconate utilisation by the human pathogen *Pseudomonas aeruginosa* requires uptake via the IctPQM TRAP transporter

**DOI:** 10.1101/2025.04.01.646390

**Authors:** Javeria Mehboob, Reyme Herman, Rory C. Elston, Heritage Afolabi, Bethan E. Kinniment-Williams, Marjan van der Woude, Anthony J. Wilkinson, Gavin H. Thomas

## Abstract

*Pseudomonas aeruginosa* PA01 is one of the major causes of disease persistence and mortality in patients with lung pathologies, relying on various host metabolites as carbon and energy sources for growth. The *ict-ich-ccl* operon (*pa0878, pa0882, pa0883*) in PAO1 is required for growth on the host molecule itaconate, a C_5_-dicarboxylate. However, it is not known how itaconate is taken up into *P. aeruginosa*. Here we demonstrate that a genetically linked tripartite ATP-independent periplasmic (TRAP) transporter (*pa0884-pa0886*), which is homologous to the known C_4_-dicarboxylate binding TRAP system, is essential for growth on itaconate, but not for the closely related C_4_-dicarboxylate succinate. Using tryptophan fluorescence spectroscopy we demonstrate that the substrate binding protein, IctP (PA0884), binds itaconate, but still retains higher affinity for the related C4-dicarboxylates. The structures of IctP bound to itaconate (1.80 Å) and succinate (1.75 Å), revealed an enclosed ligand binding pocket with ion pairing interactions with the ligand carboxylates. The C2 methylene group that is the distinguishing feature of itaconate compared to succinate is accommodated by a unique change in the IctP binding site from a Leu to Val, which distinguishes it from closely related C4-dicarboxylate binding SBPs. Together these data suggest that this transporter, which we name IctPQM, has duplicated from a canonical C4-dicarboxylate transporter and its evolution towards itaconate specificity enables this pathogen to now access a key metabolite for persistence in the host.

## Introduction

*Pseudomonas aeruginosa* (*P. aeruginosa*) is a Gram-negative biofilm forming opportunistic pathogen, which is one of the major causes of pulmonary pathologies, pneumonia, bacteraemia and other nosocomial infections [1]. This species accounts for around 7% of nosocomial infections [2] [3], and due to increasing antimicrobial resistance is classified among the WHO priority pathogens [4]. *P. aeruginosa* has been linked with high morbidity and mortality in patients suffering from chronic infections that exhibit bacterial persistence such as idiopathic pulmonary fibrosis (IPF) [5], cystic fibrosis (CF) [6], asthma [7], and chronic obstructive pulmonary disease (COPD) [8]. During infection, *P. aeruginosa* adapts to the environment of the host by regulating the expression of virulence factors such as lipopolysaccharide (LPS), alginates PsI and Pel, outer-membrane proteins and its secretion system for flagella and pili. These factors allow the bacterial cell to survive by adhesion and colonisation of host cells and contribute to chronic infection [9] [10].

In addition to cell surface factors, bacteria able to compete and colonise niches like the CF lung need to access carbon and energy sources to facilitate growth. *P. aeruginosa* has been known for a long time to prefer short-chain dicarboxylates over other more commonly favoured carbon and energy sources like glucose, and possesses a regulatory cascade that reflects this [11]. Colonisation by pathogens of other human niches like the gut relies on efficient use of host and dietary carbohydrates such as glucose [12], fucose [13] [14] and sialic acid [15], while the regulatory hierarchy of *P. aeruginosa* suggests that this is not a niche in which it would be competitive. The particular ability of *P. aeruginosa* to be a regular extracellular coloniser of the CF-lung is not totally understood. Emerging evidence suggests that the loss of the CF transmembrane conductance regulator (CFTR) results in wider metabolic dysregulation and secretion of the C_4_-dicarboxylate succinate leading to longer-term infections [16]. Interestingly, when *P. aeruginosa* consumes succinate this then triggers the host to release the related compound itaconate; 2-methylenesuccinate [16], which facilitates the characteristic chronic infection [17]. While succinate can be directly incorporated into central metabolism, the more unusual C_5_-dicarboxylate itaconate requires the products of the *ict-ich-ccl* genes (PA0878, PA0882, PA0883) to catabolise it to pyruvate and acetyl-CoA [18] [19].

Bringing these negatively charged metabolites into PA01 requires active transport by two transporters, DctA and DctPQM, which are involved in uptake of C_4_-dicarboxylates [20]. They appear to provide the only route for uptake of malate and fumarate, while residual uptake of succinate suggests an additional system(s) is present [20]. In addition to DctPQM, *P. aeruginosa* PAO1 contains 3 other DctP-family tripartite ATP-independent periplasmic transporters (TRAP) [21], two of which are phylogenetically closely related to DctPQM (PA5167-9), the closest being that encoded by the *pa0884-6* genes [21]. This system is the uncharacterised TRAP transporter that is encoded immediately downstream of the itaconate catabolic genes, with PA0884 being the substrate binding protein (SBP) and PA0885-6 the small and large membrane subunits, respectively [21]. As part of work on the catabolic enzymes in *P. aeruginosa* strain PA14, the *pa0886* gene was disrupted and growth on itaconate was reduced compared to the wild type [19], which combined with its location within the itaconate utilisation operon in PAO1 strongly suggests a function in itaconate transport [19], [20, 22]. Furthermore, introduction of the whole region including the transporter and the associated itaconate catabolic genes into the related bacterium *P. putida*, allowed the latter to grow on itaconate, supporting the idea that PA0884-6 is an itaconate-specific TRAP transporter [23]. In this study we combine genetic, biochemical and structural approaches to characterise this transporter, which we call IctPQM following the regular naming pattern for TRAP transporters, and discover a how *P. aeruginosa* has evolved a new transporter through duplication and specialisation towards itaconate over other C_4_-dicarboxylates, which enable it to utilise a key host molecule.

## Results

### The PA0884-6 TRAP transporter is essential for growth on itaconate

To investigate whether the PA0884-6 TRAP transporter has a role in the physiological uptake of itaconate, we measured growth of the wild-type strain and isogenic mutants disrupted in *pa0884* (the SBP) and *pa0885* (small membrane subunit) in minimal media supplemented with 5 mM of different carbon and energy sources (**Fig. 1**). As controls we also included a strain from the same mutant library with a disruption in an unrelated gene (*pa3728*) and a strain with a disruption in a gene encoding an essential component of the itaconate catabolic cluster (*pa0878*) [22]. All strains grow well with glucose, although the *pa0878* mutant appears to have a small growth reduction which is similar to the phenotype observed in a previous study [19]. All strains grow well on succinate and fumarate, with more rapid initial growth than with glucose, reflecting the known preference for C_4_-dicarboxylates as carbon sources [20] [24] [25]. Importantly with itaconate as the carbon source the pattern is different, with no growth for the *pa0878* catabolic mutant as expected, but also no growth for the *pa0885* mutant in the TRAP transporter membrane domain. This is consistent with a previous disruption in *pa0886*, the large membrane subunit of the transporter [19]. However, somewhat surprisingly, the *pa0884* mutant gives an intermediate phenotype, with an increased lag and reduced growth rate (**Fig. 1D**) suggesting that there may be another closely related SBP that binds to itaconate. On the basis of this *in vivo* phenotype, we suggest, following the standard naming conventions for TRAP transporters, that this system is called the IctPQM TRAP transporter.

**Fig. 1.**
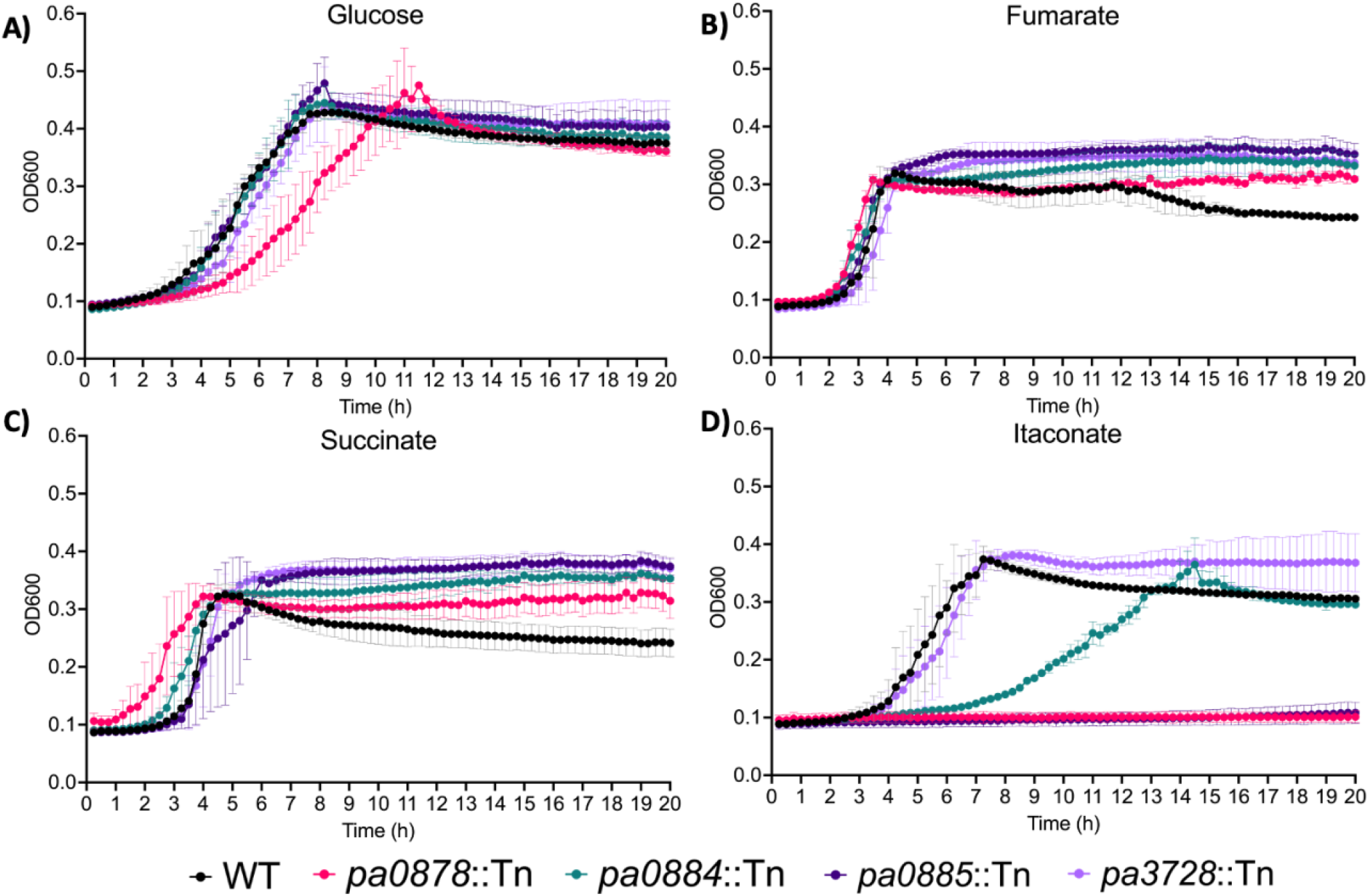
Growth curves of wildtype and mutant *P. aeruginosa* strains in minimal media containing 5mM of the carbon sources. **(A)** D-glucose, **(B)** fumarate, **(C)** succinate, **(D)** itaconate. Data were collected for 3 biological replicates.

### IctP (PA0884) can interact with C4-dicarboxylates similar in structure to succinate, including the airway metabolite itaconate

To biochemically assess the ability of the PA0884-PA0886 TRAP system in PAO1 to recognise dicarboxylates relevant to colonisation of the CF lung, we assessed their binding to recombinantly expressed IctP (PA0884) (**Supplementary Fig. 1**), the SBP of the transport system, using differential scanning fluorimetry (nanoDSF) (**Fig. 2A**). During purification, recombinantly expressed PA0884 was first partially unfolded with guanidine hydrochloride to remove any endogenous ligand likely to be present before being refolded. Upon addition of the airway metabolites succinate and itaconate to IctP, we observed an increase in thermal stability of the protein by ∼8.5°C and ∼5°C, respectively (**Fig. 2B**), whereas the sugar D-mannose gave no or very small shifts. These data provide the first biochemical suggestions that succinate and itaconate bind directly to IctP.

**Fig. 2.**
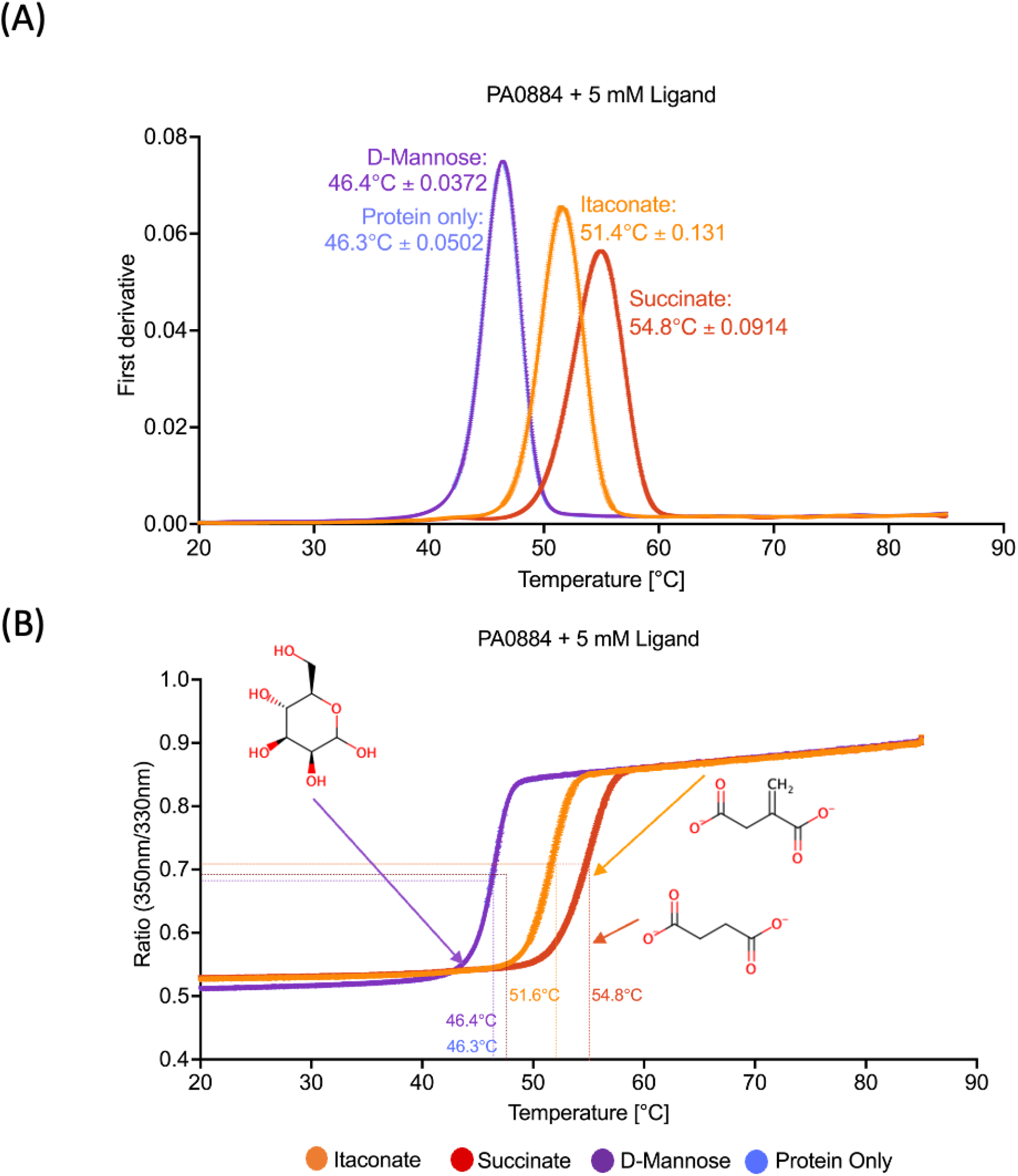
NanoDSF analysis of ligand binding to IctP (PA0884). The plots show in **A)** the first derivative and **B)** the corresponding ratio of 350nm/330nm traces in the presence of 5 mM itaconate, succinate and D-mannose. The negative control contained protein in the absence of any ligand. Structures of each ligand are included in both figures along with the *Tm* +/-SD for the three technical repeats.

### IctP can bind to itaconate but is not its preferred ligand

We next determined the binding affinities of the SBP and various ligand interactions using intrinsic fluorescence spectroscopy (**Fig. 3**). Following a similar pattern to the nanoDSF, both itaconate and succinate were bound by IctP, with respective binding affinities of 179.6 μM ± 17.1 μM and 32.6 μM ± 1.5 μM, support the finding that succinate is a tighter binder to the protein. In fact when examining binding of two other C4-dicarboxylates that are recognised by DctP-like SBPs, namely L-fumarate and L-malate, we found that L-malate exhibited tightest binding with a K_D_ of 16.2 μM ± 1.5 μM. Our biochemical analysis suggests that while PA0884 can bind itaconate, it still has retained strong binding for the regular C_4_-dicarboxylates.

**Fig. 3.**
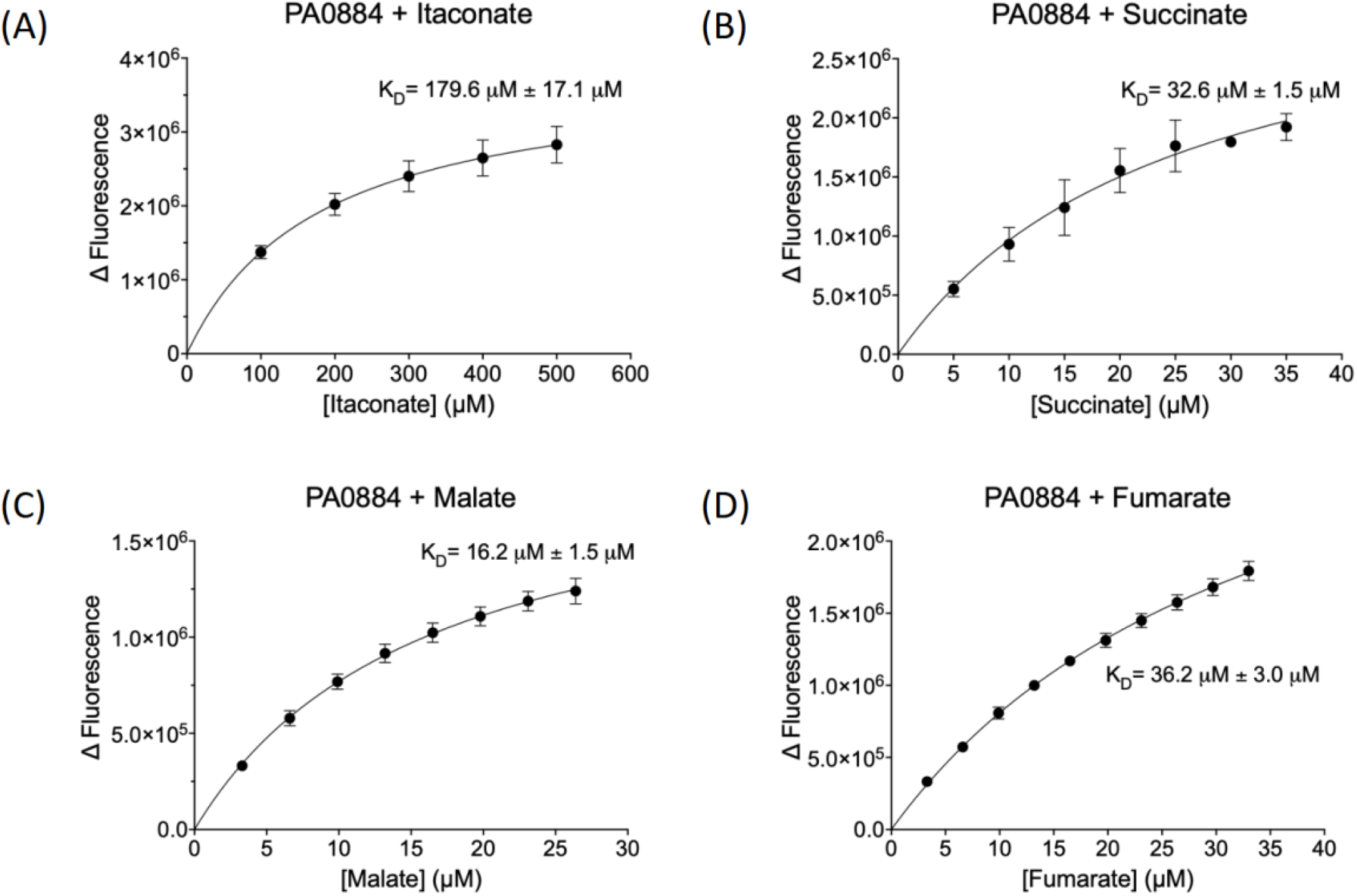
Investigation of ligand binding affinities of IctP (PA0884) to dicarboxylate ligands through measuring changes in intrinsic fluorescence. Intrinsic fluorescence was measured at 295/330nm, with ligand titrations (A) itaconate, (B) succinate, (C) malate and (D) fumarate. Standard deviations are shown for the K_D_ values.

### Itaconate is accommodated within an expanded hydrophobic pocket in the binding site of IctP

To reveal the molecular basis of binding for the dicarboxylates tested, we determined crystal structures of IctP in complexes with itaconate (PDB: 9HT3) and succinate (PDB: 9HT4). Both structures reveal the protein to be in a closed ligand-bound form similar to that seen for other TRAP SBPs [26, 27],[28],[29, 30] comprising two lobes (**Fig 4A**) with the ligands buried in an enclosed binding site. The itaconate ligand fulfils its hydrogen bonding and electrostatic potential through extensive interactions of its carboxylate groups (**Supplementary Fig 2AB**), both of which are expected to be ionised (pKa values of 3.8 and 5.5) at the pH of crystallisation. The proximal (C1) carboxylate group forms a two-pronged ion-pair with the guanidino moiety of R169, a key characteristic of a DctP family SBP ligand interaction [31, 32], further interacting with the ε-amino group of K94, the amide –NH_2_ of N209, and an ordered water molecule (**Fig. 4B**) Meanwhile, the distal (C_4_) carboxylate forms polar interactions with the charged amino group of the K41 side chain, the amide –NH_2_ moiety of the side chain of N213, the phenolic hydroxyl of Y236 and a further ordered water molecule. These polar interactions are supplemented by apolar interactions involving the side chains of V34, V35, S91 F192 and V234. The nearest neighbours of the methylene substituent are provided by the side chains of Q171, V234 and S91. This group is otherwise enclosed by the K94 side chain and a water molecule.

**Fig 4.**
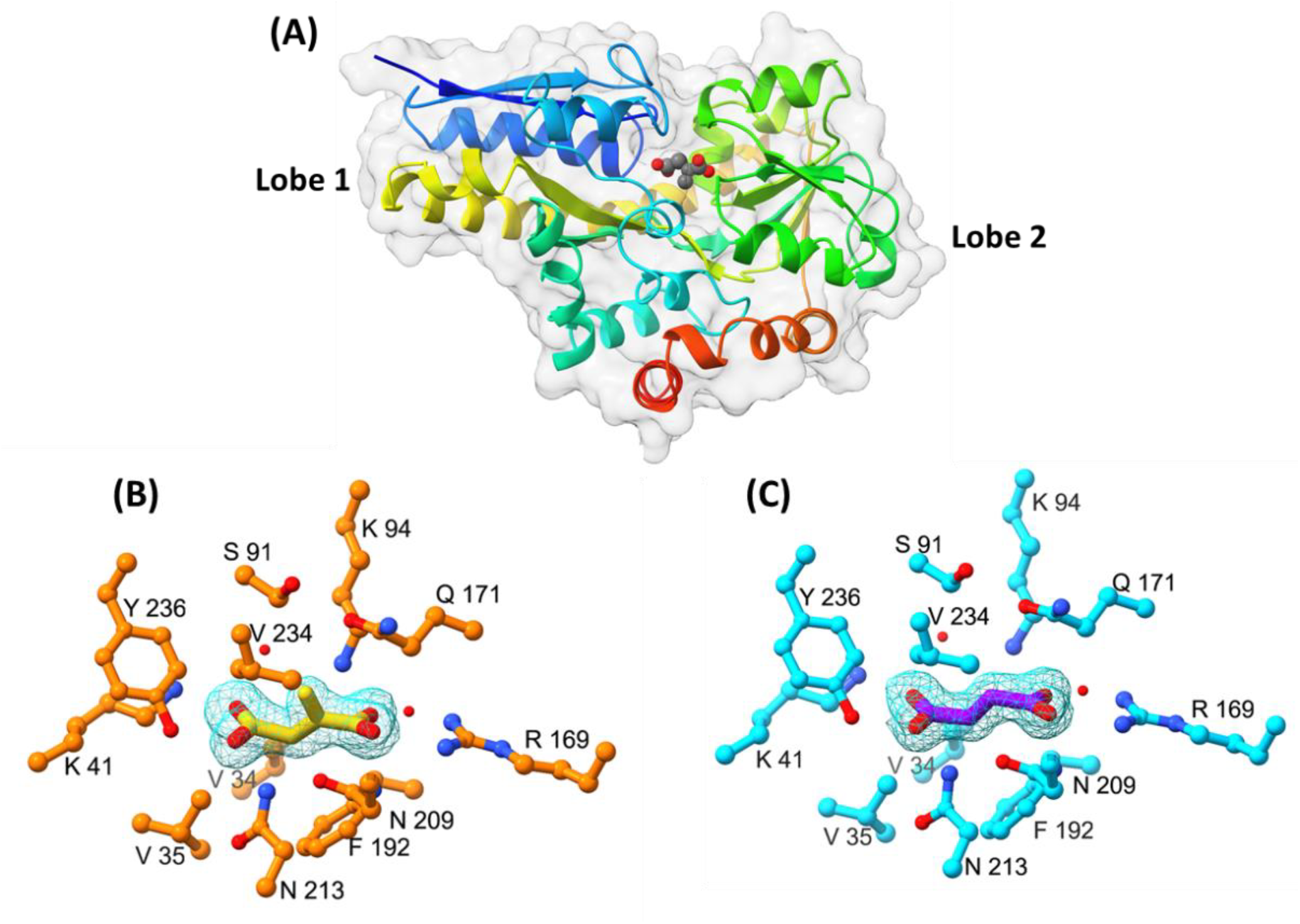
Structural characterisation of IctP with bound-ligands. **A)** Ribbon tracing of the backbone of PA0884 colour ramped from the amino terminus (blue) to the C-terminus (red). The enclosed itaconate ligand is shown in sphere format coloured by atoms with carbons in grey and oxygens in red. A transparent protein surface emphasises the burial of the ligand. **B)** Electron density map (2Fo-Fc) of the itaconate ligand (yellow) in PA0884 (orange) (contour level = 0.3σ). **C)** Electron density map (2Fo-Fc) of the succinate ligand (purple) in PA0884 (cyan) (level = 0.3).

For the succinate complex, the ligand occupies a similar volume to the itaconate and the structure of the surrounding residues is essentially unchanged (**Fig 4C**). As a result, the carboxylate groups form the same set of strong polar interactions with protein side chains as described for itaconate (**Fig 5AB**). For both ligands, the C_4_ carboxylates are directed into lobe 1 while the C1 carboxylates project towards lobe 2. The formation of these interactions would therefore be accompanied by domain closure and substrate burial as is the hallmark of ligand binding in periplasmic substrate binding proteins. The binding to the two carboxylates effectively forms a clamp around the ligand, projecting substituent on the C2 carbon down towards the long inter lobal β-strands (**Fig 4A**). We further identified a closely similar solved structure of PA0884 in complex with succinate (PDB: 9DTL), and aligned it with our structure of PA0884 bound to succinate, which has an RMSD of 0.325 Å (**Supplementary Fig 3AB**). Unsurprisingly, these two ligands bind in a highly similar manner, with V234, Q171, S91, K41, R169, K94, N209 and V34 contacts shared between them.

**Figure 5.**
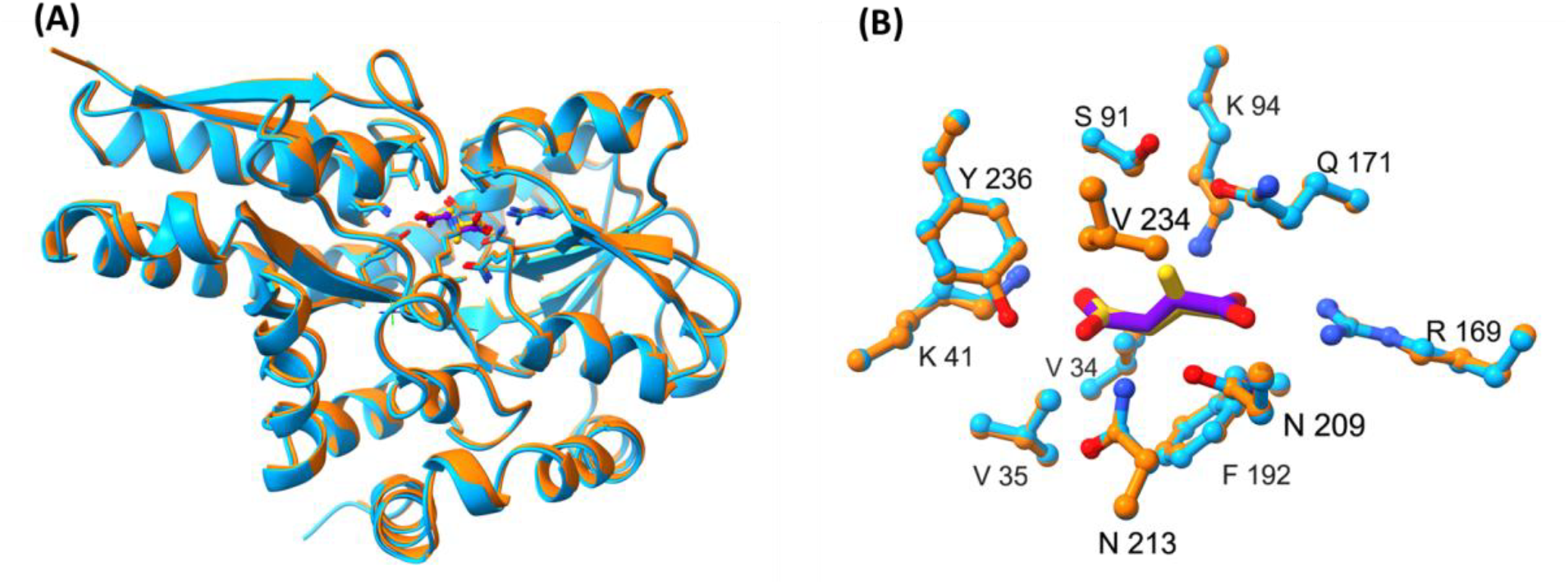
Structural alignments of PA0884 in complex with itaconate and succinate. **A)** Overlay of PA0884 bound to succinate (cyan) and PA0884 bound to itaconate (orange) **B)** A close-up view of the ligand binding site of PA0884 (cyan) bound to succinate (purple) and PA0884 (orange) bound to itaconate (yellow). Alignment algorithm used is Needleman-Wunsch and the similarity matrix is BLOSUM-62.

We then compared the structure of IctP with that of *P. aeruginosa* DctP (PA5167) bound to succinate (PDB: 9DSY) (**Fig 6**), a protein with a known function in succinate uptake in *P. aeruginosa [20]*. Secondary structure alignment of these three proteins shows absolute conservation of key binding site residues with the exception of V234 [33], reflecting the recent evolutionary relationship of these orthologous proteins **(Fig 7)**. The alteration from the larger leucine residue in DctP to the smaller valine in IctP (**Fig. 6C)**, suggests that V234 could be the primary adaptation in IctP that enables it to recognise itaconate with μM affinity. In an identical binding site, the larger L234 residue found in DctP would likely interfere with the accommodation of itaconate in the binding.

**Fig 6.**
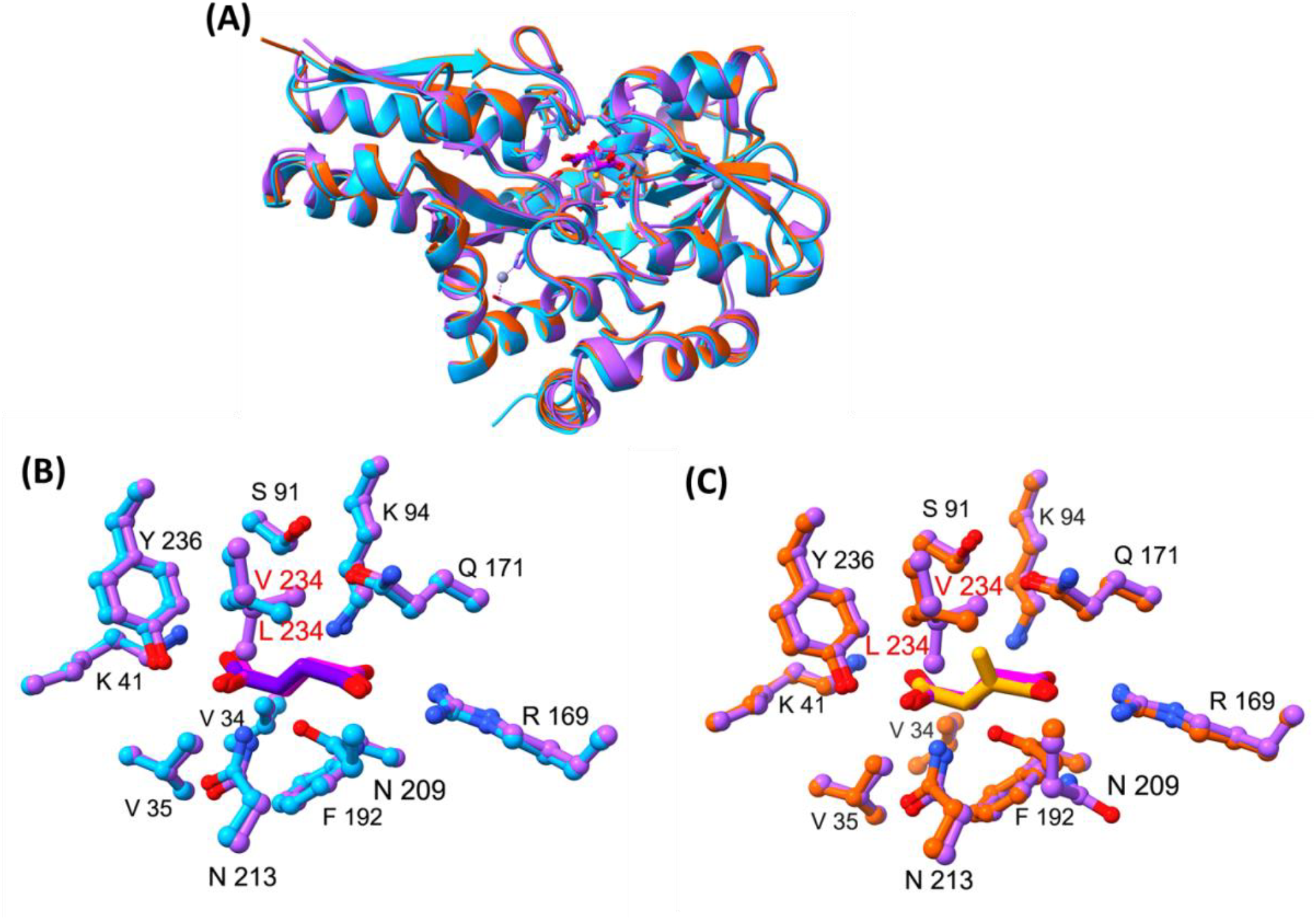
Structural comparison of IctP (PA0884) and DctP (PA5167) binding to itaconate and succinate. A) Overlay of PA0884 bound to succinate [9HT4] (cyan), PA0884 bound to itaconate (orange) [9HT3], and DctP (purple) bound to succinate (pink) [9DSY] B) A close-up view of the ligand binding site of PA0884 (cyan) bound to succinate (purple) overlayed onto PA5167 (PDB ID: 9DSY) (purple) bound to succinate (pink) C) A close-up view of the ligand binding site of PA0884 (orange) bound to itaconate (yellow) overlayed onto DctP (purple) [9DSY] bound to succinate (pink).

**Fig 7.**
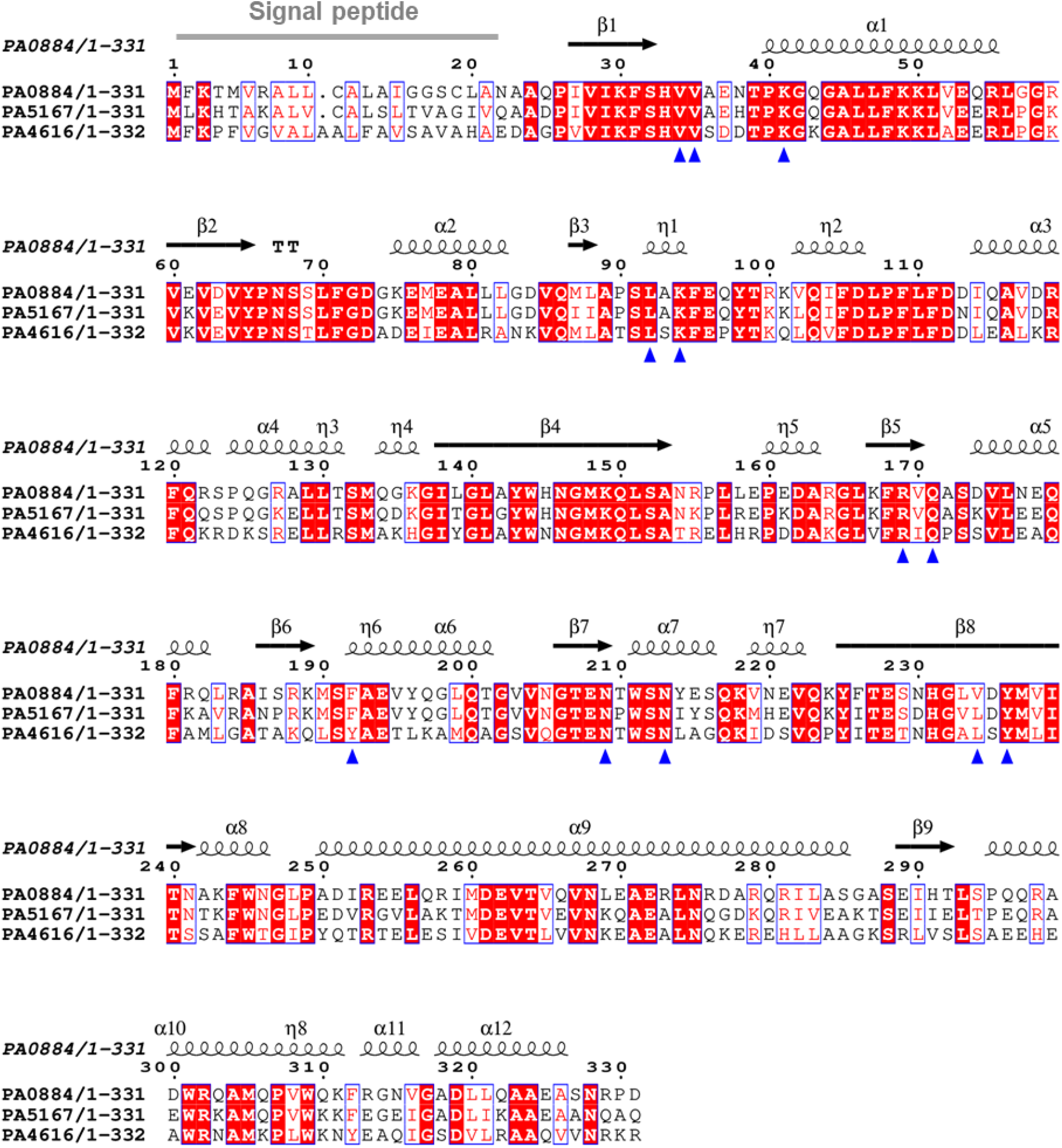
Sequence alignment of 3 PA01 TRAP SBPs with secondary structure template of PA0884 bound to succinate (9HT4). ESpript alignment of PA0884, PA5167 and PA4616 highlighting secondary structures across the sequence (created using PDB: 9HT4) and key binding site residues (blue triangles). V234 is the key binding site residue in PA0884 whereas L234 is well conserved in both PA5167 and PA4616. All the other key binding site residues are well-conserved except F192. Created with Espript.

## Discussion

The role of short-chain dicarboxylates in the establishment of chronic *P. aeruginosa* infection in the lungs of individuals with pathologies like CF and COPD has long been known. While initially being recognised as part of the immune response to infections, short-chain dicarboxylates like succinate and itaconate are now known to be used by the pathogen to reprogram itself to allow persistence in the airway [17] [34] [18]. While there is some evidence for the function of DctA and DctPQM in succinate uptake [20], how itaconate was acquired was not known. The *pa0884-pa0886* genes were suggested to encode the transporter for itaconate due to their proximity to the itaconate catabolic genes [35] and there is evidence in the form of growth assays using heterologous expression of the entire *pa0878–pa0886* cluster to support this hypothesis [23]. In this study we provide the first genetic evidence these transporter genes, which we name *ictPQM*, are essential for itaconate uptake and biochemical and structural data to suggest that IctP binds itaconate and also succinate and other dicarboxylates.

As *P. aeruginosa* PAO1 has a preference for succinate as a carbon source [36–38] it is perhaps not surprising that it can also utilise other dicarboxylates like itaconate as sole carbon source, supporting comparable growth rates to succinate [20]. While for succinate there are multiple transporters of varying affinity that can take up succinate [20], our data suggests that the TRAP transporter is the sole route of uptake in *P. aeruginosa* PAO1 (Fig. 1D). However, this genetic analysis revealed that while the small membrane component was essential for growth with itaconate, the disruption of the SBP did not completely abolish growth. This result is on the one hand a surprise, as the interactions between ‘cognate ‘SBPs and their membrane components is usually such that they are unique, although in SBP-dependent ABC transporter there are examples of multiple SBPs interacting with the same membrane domains [39]. On the other hand, given the high sequence similarity (73.6%) of the IctPQM system to the canonical DctPQM system, it is likely that the proteins are still similar enough for a ‘non-cognate ‘interaction of a highly similar SBP, namely DctP (PA5167), with IctQM could occur, which is what we suggest is happening here. This is consistent also with our biochemical and structural data which shows the highly similar binding site between DctP and IctP and that C_4_-dicarboxylate binding is still retained in IctP. Examining the distribution of the IctPQM system, its presence in only a small number of related species of *Pseudomonas* suggests clear that it has evolved via a duplication of the *dctPQM* genes in this lineage of bacteria and then subsequent specialisation towards itaconate as a substrate, although it clearly retains many of its ‘ancestral ‘binding features.

TRAP transporters are interesting hybrid transporters that combine high ligand binding affinity and specificity to concentrate substates using the membrane potential, as opposed to via direct ATP hydrolysis as used by SBP-dependent ABC transporters [40, 41]. It is interesting that while other ‘scaffolds ‘such as the DctA C_4_-dicarboxylate transporter are present in *Pseudomonas* sp., it has been the duplicated and adaptation of a TRAP system that has been selected during evolution, perhaps suggesting that the concentrations of itaconate in the environment where these more ancestral Pseudomonads evolved was in the low μM range. The duplication of the SBP to confer additional binding range to a transporter has been seen multiple times in ABC transporters [42], but here the duplication of the whole transporter and its integration into an itaconate-regulated operon [19], suggests that the bacteria has selected to genetically separate out the ancestral C_4_-dicarboxylate transport function to that of the new itaconate specific IctPQM.

It is interesting to link this observation back to the role of itaconate transport and catabolism in infection biology. As *P. aeruginosa* likely had this functioning itaconate utilisation system before its specialisation as a human pathogen, it is possible that helped enable successful host colonisation through the removal of this potentially toxic molecule. Thinking more broadly, considering that in both skin wound settings and in the CF lung the presence of *P. aeruginosa* is often found in polymicrobial CF infections [43] [44], it possible that the removal of itaconate by *P. aeruginosa* is a feature of these ecosystems. For example, *S. aureus* lung infections induce itaconate release in the airway through the activation of host *IrG1[45]*, which is known to be bactericidal to *S. aureus* [46]. However, unlike *P. aeruginosa, S. aureus* is thought to be unable to degrade itaconate [47], so in a polymicrobial infection it is possible that *P. aeruginosa* could alleviate the stress induced by itaconate for both itself and *S. aureus* by removing and degrading this molecule to allow persistence of both species in the airway.

While itaconate utilisation suggests a direct route to increased colonisation, there is also data suggesting more indirectly impact of external succinate and itaconate on the state of *P. aeruginosa* growth, with studies suggesting succinate inhibits biofilm formation, while another that itaconate promotes it [16, 18]. To see if the lack of uptake due to loss of IctPQM function altered biofilm formation, an initial experiment suggested that itaconate, but not succinate, promoted biofilm formation in the absence of the transporter (**Supplementary Fig 4**). While it is unclear what mechanisms are occurring here, it raises the possibility that itaconate is doing more than just being consumed by *P. aeruginosa* as a carbon and energy source, perhaps as a result of itaconate remaining outside the cell, for a longer period of time prolonging outer membrane stress which leads to increased biofilm formation.

In conclusion, our definition and analysis of the IctPQM TRAP transport system in this work, reveals a lineage-specific duplication and specialisation of an ancestral C_4_-dicarboxylate transporter, DctPQM, for a function in itaconate uptake and the description of the first dedicated itaconate uptake system in bacteria.

## Methods

### Cloning and protein production

The SBP domain of the transporter operon *pa0884* was cloned into pETYSBLIC3C [48] and expressed in *E. coli* BL21(DE3). Cultivations were performed in baffled 2L flasks on an orbital shaker at 120 rpm at 37°C. 1L expression cultures were set up in LB media with a starting OD (600 nm) of 0.1 from overnight cultures. The expression cultures were induced with 0.5 mM IPTG at an OD 0.6 (at 600nm). Overnight cultures were spun down and cell pellets re-suspended in a 30ml 20mM KPi + 200 mM NaCl wash, 5 mM imidazole buffer and 10% glycerol. Re-suspended cells were lysed with sonication and clarified supernatant collected by spinning down at 27000 x g for 40 mins at 4°C. Supernatant was loaded onto a 5 ml FF - HisTrap column with an isocratic elution method with 20mM KPi pH 8, 200 mM NaCl, 500 mM imidazole and 10% glycerol elution buffer using AKTA Pure (Cytiva). Ni-affinity purification elution fractions were pooled together and loaded onto Supradex S-200 column (GE Healthcare Life Sciences Buckinghamshire, UK) for buffer exchange into SEC buffer (20mM Tris-HCl pH 8, 50mM NaCl).

### Protein melting temperature determination

NanoDSF was performed using Prometheus NanoTemper technologies equipped with a back reflection mode (NanoTemper Technologies, Munchen, Germany). 10 μl samples were loaded into the nanoDSF capillaries (NanoTemper Technologies, München, Germany) and placed in the sample holder. A final concentration of 5 μM protein in 50 mM KPi, 200 mM NaCl and each ligand at final concentrations of 5 mM was analysed in triplicates. Samples were heated up at a thermal ramping rate of 1°C/min from 25 to 95°C and intrinsic protein fluorescence was measured from 330 and 350 nm with a dual detector. The excitation power for the different samples varied, but was sufficient for A330 and A350 fluorescent signals to exceed 2000 relative fluorescence units (RFU). Protein melting temperature (*T*_m_) and aggregation (*T*_*agg*_) were also calculated from the data and plotted in GraphPad (Prism 10).

### Fluorimetry

Intrinsic tryptophan fluorescence was measured with the FluoroMax 4 fluorescence spectrometer (Horiba Jobin-Yvon) at room temperature. PA0884 was prepared in 50 mM KPi pH 7.8 with 200 mM NaCl buffer. The protein was used at a concentration of 0.5 μM and excited at 295 nm with various ligands added at 100 μM. The slit width was 3.5 nm and the emission spectra was collected at 330 nm. Protein samples were titrated with increasing concentrations of ligands fumarate, malate, succinate and itaconate. *K*_D_ was calculated from the hyperbolic fit of the binding curve and plotted using GraphPad (Prism 10).

### Crystallisation of PA0884-succinate and PA0884-itaconate

For crystallisation, recombinantly expressed PA0884 was concentrated to 15 mg/mL in 20 mM Tris-HCl pH 8, 50 mM NaCl and mixed with either 50 mM itaconate, pH 7 or 50 mM succinate, pH 7. The mixture was incubated in solution at room temperature for 15 minutes. Protein crystals were grown in sitting drops (1:1 ratio) over the crystallisation solution (JCSG-plus HT-96 D12: 0.04 M Potassium phosphate monobasic, 16 % w/v PEG 8000) at room temperature in MRC 96-well 2-drop crystallisation plates. Crystals of both protein-ligand complexes were cryo-protected using 20% glycerol.

### Data collection and structure determination

X-ray diffraction data were collected at the Diamond Light Source, UK on Beamline i03 for crystals of both PA0884-succinate and PA0884-itaconate. The data collected for both protein crystals were indexed and scaled using the DIALS pipeline [49]. Data reduction was performed using AIMLESS 0.7.13 [50]. The molecular replacement program MOLREP [51] was used to obtain initial phase information for both protein crystals using the predicted structure of PA0884 generated using AlphaFold (AF-Q9I561-F1). The respective structures were refined using REFMAC5 [52]. Refined coordinate sets and structure factors were deposited into the PDB with the entry codes 9HT3 for PA0884-itaconate and 9HT4 for PA0884-succinate. Data collection statistics are provided in **Supplementary Data Table S1**.

### Structural analysis

The buried surface area between the protein and ligand was calculated using the ‘measure buried area ‘command in ChimeraX 1.8 [53]. The binding site was defined as atoms within 5 Å from the ligand. Solvent accessible surfaces were determined using a probe radius of 1.4 Å which is used to approximate a water molecule. As a result, the buried surface area between the surface of the ligand and the surface of the surrounding protein residues could be determined.

### Growth data

Strains used in the growth assays have the following genotypes: WT strain PAO1, enzymatic pathway mutant pa0878-E03::ISlacZ/hah, substrate binding protein domain pa0884-F06::ISlacZ/hah, small subunit of the membrane domain pa0885-C09::ISlacZ/hah and a gene of unrelated function as a negative control pa3728-C01::ISlacZ/hah. WT and mutant strains were grown overnight in LB media. Overnight cells were pelleted and washed with MilliQ water 1000 x g at 4 °C. Cells were resuspended in Neidhart MOPS M9 minimal media with carbon sources itaconate, succinate, fumarate and D-glucose. Inoculation culture was added to Nunc™ MicroWell™ 96-Well Microplate wells to M9 minimal media and carbon sources.

## Conflict of interest

All other authors declare that they have no conflicts of interest with the contents of this article.

## Acknowledgments

This work was supported by the Biotechnology and Biological Sciences Research Council grant BB/W510373/1 supporting J.M., BB/W510531/1 supporting R. H, BB/T017805/1 for use of the X-ray facility and BB/X003035/1 for general support of the GHT lab.

We thank Johan Turkenburg and Sam Hart for help with data collection and staff at the DIAMOND light Source at Harwell for excellent X-ray facilities. We would also thank Dr Andrew Leech in the Technology Facility, Department of Biology, University of York for help with interpretation of data from nanoDSF and fluorimetry.

## Author contributions

R. H., and G. H. T. conceptualization; J. M., R. H., R. E., and G. H. T. validation; J. M., R. H., B. K. W., R. E., A. J. W., and G. H. T. formal analysis; J. M., R. H., R. E., and G. H. T. investigation; J. M., R. H., B. K. W., R. E., H. A., A. J. W., and G. H. T. data curation;. G. H. T. resources; J. M., R. H. writing–original draft; J. M., R. H., B. K. W., R. E., A. J. W., and G. H. T. writing–review & editing; J. M., R. H., B. K. W., and A. J. W. visualisation; A. J. W., M vd W, and G. H. T. supervision; R. H., J. M., R. E., and G. H. T. project administration; A. J. W., and G. H. T. funding acquisition.

## Notes

https://www.rcsb.org/structure/9HT4

https://www.rcsb.org/structure/9HT3

### Competing Interest Statement

The authors have declared no competing interest.

