## Supplementary data for "Itaconate utilisation by the human pathogen *Pseudomonas aeruginosa* requires uptake via the IctPQM TRAP transporter"

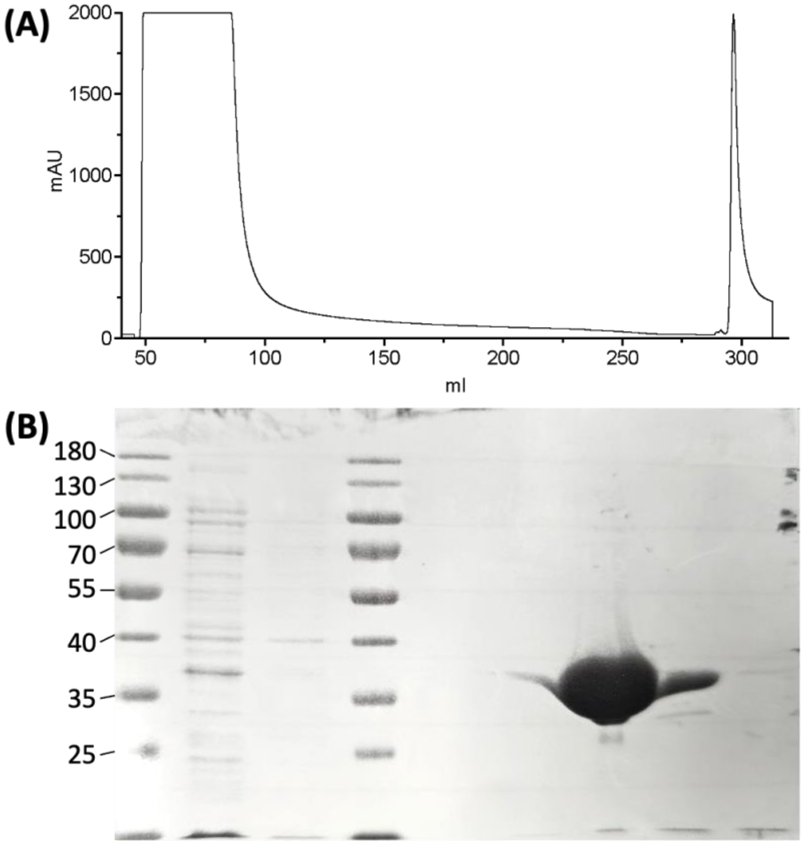


**Supplementary Fig 1 (A)** Purification trace of PA0884 **(B)** Coomassie gel of the flow through, wash through and elution fractions.

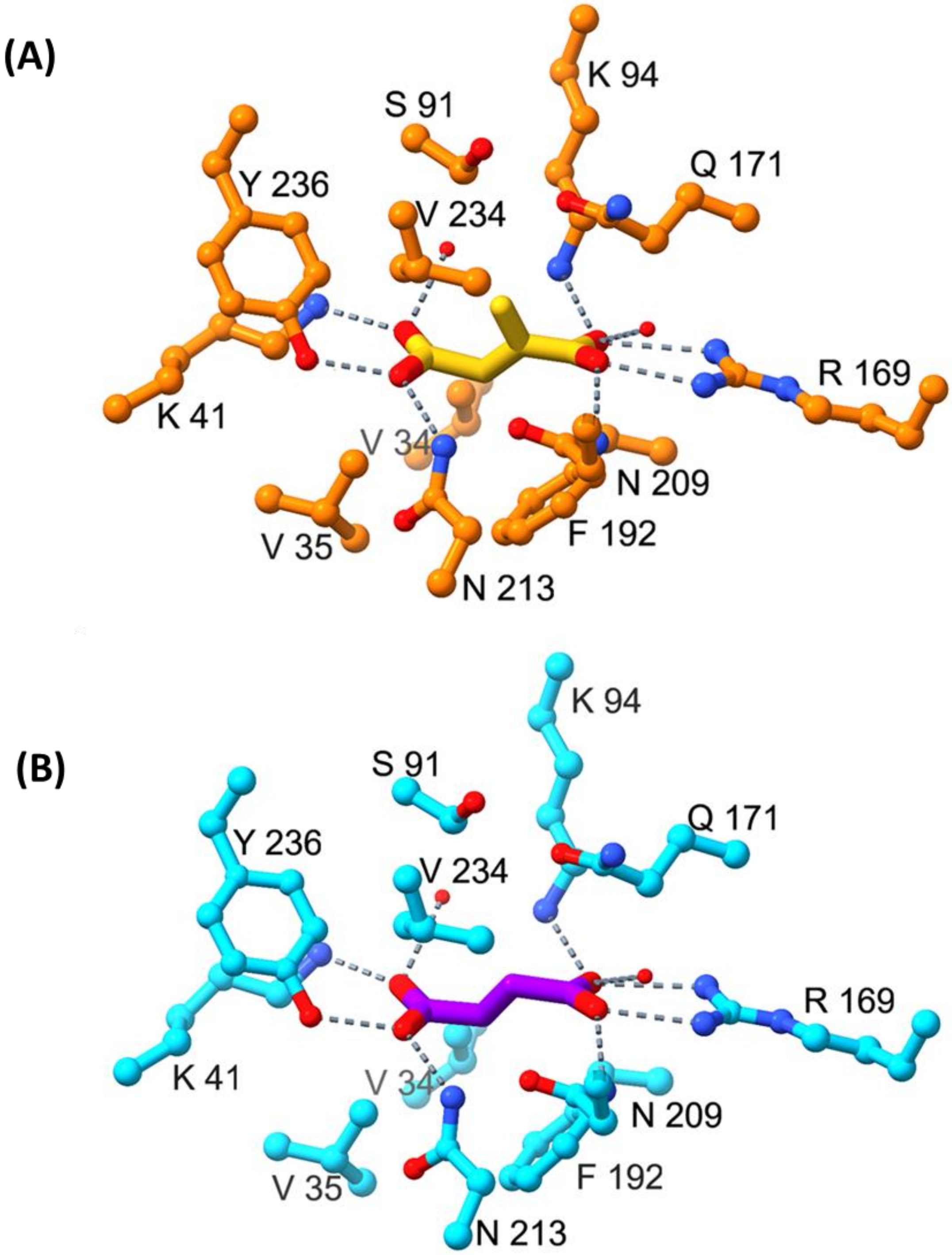


**Supplementary Figure 2.** Hydrogen bonding and electrostatic interactions of PA0884 in complex with **(A)** itaconate (yellow) and **(B)** succinate (purple)

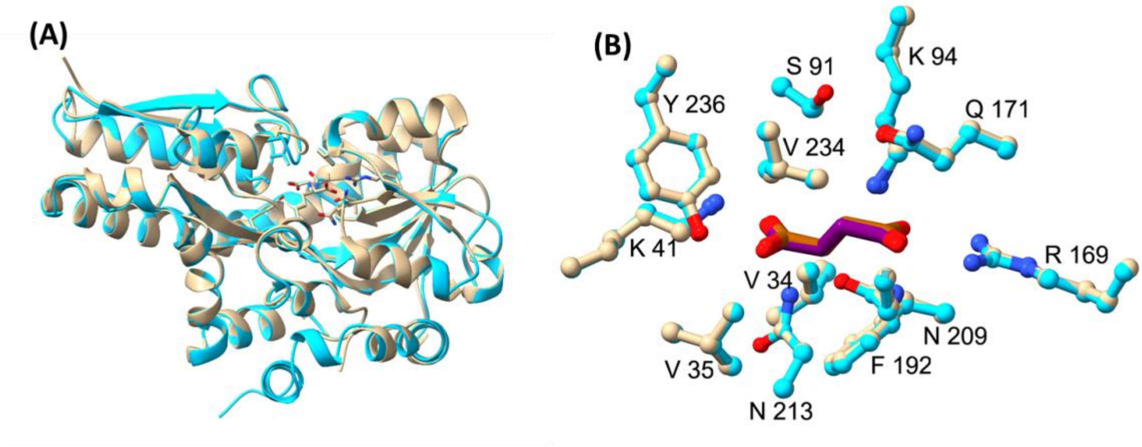


**Supplementary Figure 3 A)** Overlay of PA0884 bound to succinate (cyan) and PA0884 (9DTL) bound to succinate (beige) **B)** A close-up view of the ligand binding site of PA0884 (cyan) bound to succinate (purple) and 9DTL (beige) bound to succinate (brown). This demonstrates the structures are nearly identical but 9DTL is missing some of the C terminal end that our structure has. (stops at Arg313 while our structure continues to Asp331).

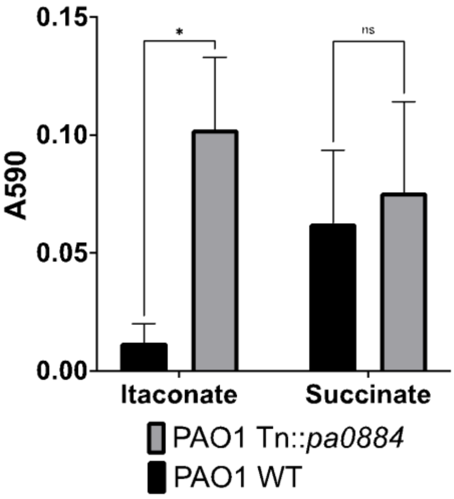


**Supplementary Figure 4**. Biofilm assay of the WT and *Tn::pa0884* mutant in itaconate and succinate. *P. aeruginosa* strains were grown overnight in LB broth at 37 degrees and washed three times with 1X M9-salts (3 g $L^{-1}$ $KH_2PO_4$, 0.5 g $L^{-1}$ NaCl, 6.78 g $L^{-1}$ $Na_2HPO_4$, 1 g $L^{-1}$ $NH_4Cl$). The washed cells were resuspended in either M9-succinate (1X M9-salts, 1 mM MgSO4, 1 mg $mL^{-1}$ Thiamine-HCl, 50 mM succinate) or M9-itaconate (1X M9-salts, 1 mM MgSO4, 1 mg $mL^{-1}$ Thiamine-HCl, 50 mM itaconate). The cells were used to inoculate fresh M9 media to an OD600 of 0.05 in a 96-well polystyrene plate. Biofilms were allowed to grow static at 37°C for 24 hours. The resulting biofilms were then stained with crystal violet and quantified by measuring the absorbance at 590 nm.

| | PA0884-itaconate | PA0884-succinate |
|---|---|---|
| PDB ID | 9HT3 | 9HT4 |
| Source Beamline | Diamond Beamline i03 | Diamond Beamline i03 |
| Wavelength | 0.9763 | 0.9763 |
| Type | Synchrotron | Synchrotron |
| Detector | Dectris EIGER2 XE 16M | Dectris EIGER2 XE 16M |
| **Data statistics** | | |
| Space group | $P2_12_12_1$ | $P2_12_12_1$ |
| Cell constants a, b, c (Å) | 34.206, 117.490, 166.063 | 33.957, 118.526, 167.210 |
| Cell constants α, β, γ ( ◦ ) | 90.000, 90.000, 90.000 | 90.000, 90.000, 90.000 |
| Overall resolution range (Å) | 50.07 – 1.80 | 50.44 – 1.75 |
| Inner shell resolution range (Å) | 50.07 – 9.00 | 50.44 – 9.09 |
| Outer shell resolution range (Å) | 1.84 – 1.80 | 1.78 – 1.75 |
| Overall completeness (%) | 100 | 100 |
| Rmerge for all I+ and I-[a] | 0.317 | 0.415 |
| < I/σ(I) > | 7.1 | 5.3 |
| Overall multiplicity | 13.1 | 13.0 |
| **Refinement Statistics** | | |
| Program | REFMAC | REFMAC |
| Number of observations | 830168 | 907802 |
| Number of unique reflections | 63463 | 69593 |
| R, Rfree [b,c] | 0.18, 0.23 | 0.20, 0.24 |
| Average B, all atoms ($A^2$) | 24.75 | 24.944 |
| RMS Deviations – Bonds[d] | 0.0153 | 0.0154 |
| RMS Deviations – Angles[d] | 2.575 | 2.553 |
| Ramachandran outliers (%) | 0 | 0 |

**Supplementary Table S1.** Data collection statistics for PA0884 crystal structures bound with itaconate and succinate.

[a] Rmerge = ∑H K L∑I |/I - */∑H K L∑I where /I is the intensity of the ith measurement of a reflection with indexes hkl and I is the STATISTICALLY WEIGHTED AVERAGE REFLECTION INTENSITY.*

[b] *R-factor =||Fo| −|Fo||/|Fo| where Fo and Fc are the observed and calculated structure factor amplitudes, respectively.*

[c] *R-free is the R-factor calculated with 5 % of the reflections chosen at random and omitted from refinement.*

[d] *Root-mean-square deviation of bond lengths and bond angles from ideal geometry.*
